# Spawning Biology and Induced Breeding of freshwater catfish *Mystus dibrugarensis*: An approach to conservation

**DOI:** 10.1101/229377

**Authors:** Bhenila Bailung, S. P. Biswas

**Affiliations:** Life Sciences department, Dibrugarh University, Assam, India

**Keywords:** bagridae, breeding, reproductive aspects, spawning

## Abstract

Spawning biology of hormone induced bagridae catfish, *Mystus dibrugarensis* was observed in captive condition. Active involvement of male in courtship after 5-9 hr. of hormone administration was observed. Different doses of ovaprim 0.4, 0.6, 0.8, 1.0 ml/kg were used by maintaining 1:1, 2:1 and 3:1 ratio of M:F. All female successfully bred in all the condition in more or less amount. Overall 24.5-77.5% fertilization and 12-76% hatching rate was observed. However, considering the rate of fertilization, and hatching success in the present case, at 0.8 ml/kg body weight and sex ratio 2♂: 1 ♀ seems ideal for induced breeding of *M. dibrugarensis*. Apart from induced breeding certain reproductive aspects were also studied in the present experiment.

## INTRODUCTION

North East India is a huge reservoir of fresh water fish germplasm and considered as a hot spot of fresh water fish biodiversity (Mahanta *et al.,* 2001). Over 200 fish species have been reported from the Brahmaputra drainage system so far (Sen, 2000), of which 78 species have been identified as highly suitable for aquarium rearing (Biswas *et al.,* 2007). Due to various factors like over-exploitation, habitat destruction these species facing lots of threat as a result there no. decreases day by day. *Mystus dibrugarensis* (Chaudhuri, 1913) is a freshwater riverine catfish having ornamental value is one of them. According to IUCN it is considered under least concern, but unfortunately it is found in certain pockets of upper Brahmaputra basin mainly in Dihing River (Bailung, 2014). Captive breeding of fish, is a widely used management tool in attempts to restore and conservation of the wild populations of endangered and endemic species and simultaneously to supplement and enhance yields for fisheries (Fleming, 1994). Considering the potential of fish species, captive breeding can serve the purpose for introducing the indigenous species in the global market generating foreign exchange (Das and Kalita, 2003). Therefore, to save species from its extinction through induced breeding a complete knowledge about its biology including breeding behaviour, fecundity, fertilization and hatching through captive breeding is also essential (Islam *et al,* 2011). Successful technology for breeding and rearing of a species in captivity is the prerequisite for rehabilitation of natural stocks as well as for culture. Therefore, standardization of breeding protocol suitable to the climate condition of Assam will definitely boost the entrepreneurs of native fish species.

Different workers worked on spawning season and induced breeding of *Mystus* species such as Ray, 2005 (*Mystus gulio*); Islam *et al,* 2011 (*Mystus vittatus*); Gupta *et al.,* 2013 (*Mystus tengara*). But till date there is no any information about induced breeding of *Mystus dibrugarensis*. In view of the paucity of knowledge on the breeding practice and its occasional occurrence, *Mystus dibrugarensis* is selected for a detailed study.

## METHODS & METHODOLOGY

Live and healthy species *M. dibrugarensis* were collected from Dihing River of Dibrugarh district, Assam, India from its confluence with almighty R. Brahmaputra i.e. Dihingmukh (27°15^/^41^//^ N and 94°42^/^10^//^ E) of upper Assam from January 2014 to August 2016.

Sex differentiation of *M. dibrugarensis* was done by careful visual inspection of the presence of genital papilla and genital pore. Sex-ratio was determined by separation of specimen from the monthly collection into male and female groups and counting the total number of specimen in two sex groups by following Mahmood *et al.* (2011). Chi-square test (χ^2^) was done to test the ratio difference was significant or not, assuming that the ratio of male to female in the population to be 1:1. Routine assessment of gonadal development is normally done by assessing individuals to stages by characters which can be differentiated with the naked eye. Depending on the morphology of gonads and portion of abdominal cavity occupied by gonads, both testicular and ovarian cycle of male and female respectively were done by following Nikolsky (1963).

Ovaries from mature specimens were only considered for fecundity studies. Sub-sampling method (Bagenal, 1957a) was employed for calculating fecundity. Likewise relative fecundity was estimated by dividing the absolute fecundity with total body weight (Biswas, 2002). Ova-diameter measurement was done with the help of stereo microscope and Leica software.

Breeding trails were done from late April to July of three consecutive years (2014-2016). The collected specimens were subjected through a 30-day quarantine period as per (Louis, 1995). Brooders were stocked in separate earthen ponds (3×3×1 m) having sandy bottom from January to March. In April, specimen were transferred to fibre circular tank (135D×35H cm) and were kept there until breeding experiment was conducted. Tanks were provided good quality of water, with the facility of water circulation, aeration and replenishment. A mixture of sand and small gravel of about 4cm was filled at the bottom and some aquatic plants like water hyacinth (*Eichhornia crassipes*) and *Pistia* was placed to create a spawning habitat. They were supplied with natural food like live earthworm, mosquito larvae, small insect, ants, and plankton as well as artificially formulated food such as dried *Tubifex*, prawns, *Artemia* and fishes twice daily at 5% of their body weight. The physico-chemical parameters of water was determined as per APHA (1998) and digital gadgets.

## RESULTS & DISCUSSION

Sex differentiation in *M. dibrugarensis* is relatively simple. Male having a soft conical projection in front of the anal fin that termed as genital papilla. In female this genital papilla was absent whereas they possess a round genital opening and swelling abdomen when they were matured. This observation was also recorded by various workers in other *Mystus* species (Bhatt, 1971a and b; Ng, 2001; Musa and Bhuiyan, 2007; Darshan *et al,* 2011 and 2013). Sex-ratio provides information on the proportion of male and female fish in the population. The overall sex-ratio was 0.932:1 (M: F), significantly tilted towards female (*χ*^2^ = 5.87, at P _0.01_). Here, out of total specimens examined, 45.7% was males and 54.2% was female. Monthly distribution of sexes fluctuated significantly in favor of female in April, May, June, July is may be because of their heavier weight due to their full gonadal development (Biswas, 1984). This observation was similar with the Bhatt (1971a); Rao and Sharma (1984). Hypoxia or low dissolved oxygen levels have been shown to favors male dominance in populations due to the effect on sexual development and sex differentiation (Shang *et al.* 2006). This may be the reason of male dominance during winter months.

Depending on the morphology of testis and ovary, portion of abdominal cavity occupied by gonads, at different developmental period and size of the intra-ovarian oocytes, five maturity stages of gonad have been identified (Nikolsky, 1963) namely stage I: Immature; stage II: maturing; stage III: Mature, stage IV: Ripe and stage V: Spent. Gupta & Banerjee (2013) and Basu *et al.* (2015) mentioned five maturity in *Mystus tengara* and *M. vittatus.* The gonads in immature stage is highly rudimentary and pinkish in colour. In maturing stage gonads occupied about half of the body cavity and vessels started to developed. Gonads in mature stages occupied 3/4^th^ of the body cavity with well-developed blood vessels. Eggs were also clearly seen in ovary through naked eyes. As the maturity progressed, the gonads became thicker, changed to yellowish or reddish in color due to development of blood vessels and it occupied almost the entire ventral cavity. Gonads became highly reduced in size and flaccid in spent stage. Here, development of finger like projection was observed in testes from maturing stage which was found gradually increased in size and volume as the maturity progressed. Gupta and Banerjee (2013) called this projection as testicular in *M. tengara.* Ripe females were found from April to August with a peak in June whereas ripe males were observed from March to August with the highest percentage encountered in May. Monthly percentage distribution of females and males at various maturity stages are given in Table 1.

**Table 1:**
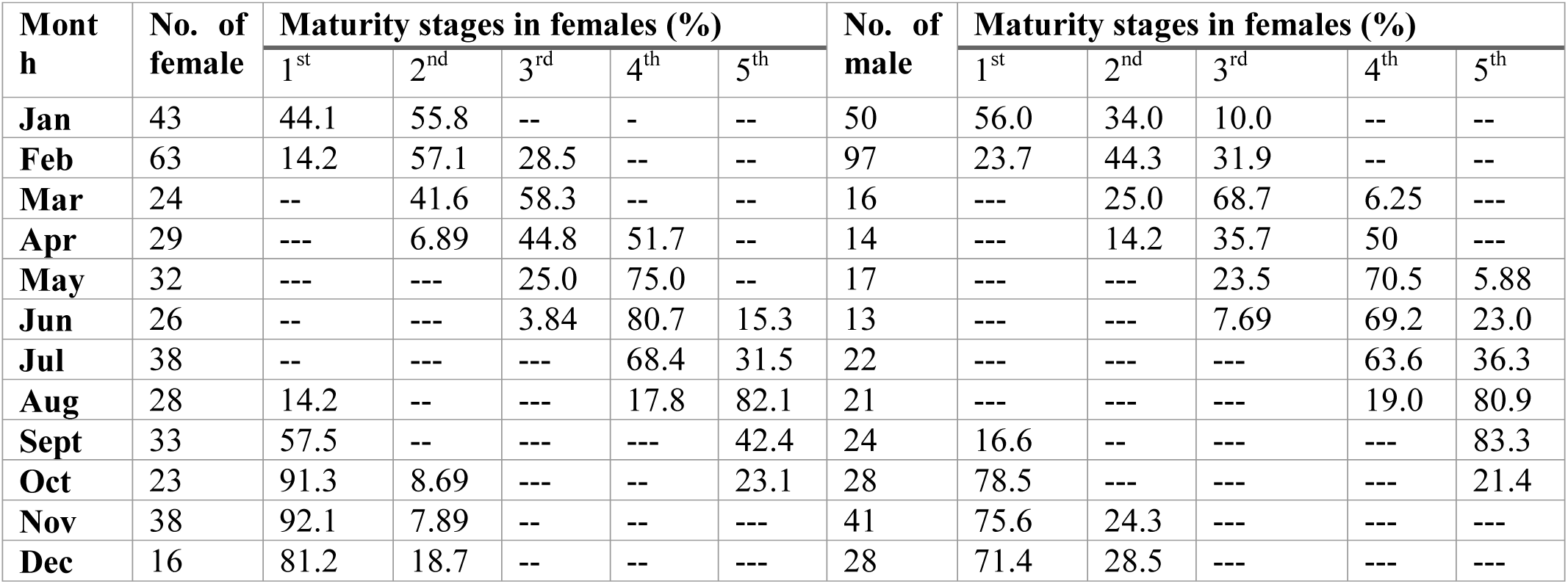
Monthly occurrence of female in different stages of maturation

The development of gonad can be represented by an index called gonadosomatic ratio (GSR) which provides information about spawning season of a particular species. Progressive increase of GSR value in both the sexes from April indicated that its breeding period starts from April. High GSR value in female was recorded in (8.15±3.27) April, (12.4±3.8) May, (11.1±4.06) June and (6.58±0.68) July. Similarly, high GSR in male was (1.21±0.73) April, (1.66±0.41) May, (1.49±0.45) June and (1.27 ±0.44) July. The ova diameter of *M. dibrugarensis* progressively increased from January onwards and maximum size of intra-ovarian egg was observed in May (1.28±0.24) to July (1.31± 0.17). This suggested that these months as peak breeding period of the selected species. Declination in the GSR value from August indicated that the spawning is over by August in *M. dibrugarensis.* This result was similar with the finding of Sarker *et al.* (2002) and Basu *et al.* (2015).

Fecundity provides information about reproductive potential of the spawning stock. The maximum absolute fecundity (12338.57 ± 1241.04) was found in the length group of 11.5-14 cm and the minimum (6965±889.38) in 6.5-9.0 cm. The fecundity was found increased with the increase in length of the *M. dibrugarensis* that similar with the findings of Azadi *et al.* (1987), Siddique *et al.* (2008), Islam *et al.* (2011) and Gupta (2013). Monthly, highest absolute fecundity (7026±2200) and relative fecundity (830.2 ±167.8) was found in April and lowest absolute fecundity (411±154.7) and relative fecundity (44 ± 22.7) in August. They were categorized as single spawner because it was found that they did not release the remaining eggs which were found reabsorbed later. Similarly Prabhu (1956), Qasim and Qayyum (1961), Bhatt (1971a), and Rao and Sharma (1984) reported *M. vittatus* as single spawner, Gupta and Banerjee (2013) also reported *M. tengara* as single spawner.

For induced breeding, total length and weight of the fish were measured nearest to 1.0 mm and 0.1 g, respectively. Hormone “ovaprim” was induced below the pectoral fin with 1ml syringe at a dose of 0.4-1.0 ml/kg during evening time by maintaining 1:1, 2:1 and 3:1 ratio of M:F. After 5-9 hr. male was observed more actively involved in courtship. The same phenomenon was also reported by Alam *et al.* (2006) and Islam *et al.* (2011). Spawning occurred after 2-3 hr. of latency period and continued for 1-2 hr (Bailung and Biswas, 2014). Eggs adhesive, (0.9±0.2) mm in size, creamy or yellowish in color. Fertilization rate 24.4%-55.6%; 30%-77.5% and 26.5%-65.7% and hatching rate 20-66%; 22-76% and 12-58% was observed in 1:1, 2:1 and 3:1 ratio of M:F respectively. Likewise, overall 20.4-54.3%; 24.4-67.06; 31.1-77.54 and 26.5-70.3% fertilization was observed in 0.4, 0.6, 0.8 and 1.0 ml/kg hormone dose. However, considering the rate of fertilization, and hatching success in the present case, at 0.8 ml/kg body weight and sex ratio 2♂: 1 ♀ seems ideal for induced breeding of *M. dibrugarensis.* Similar finding was reported by Islam *et al.* (2011).

**Table 2:**
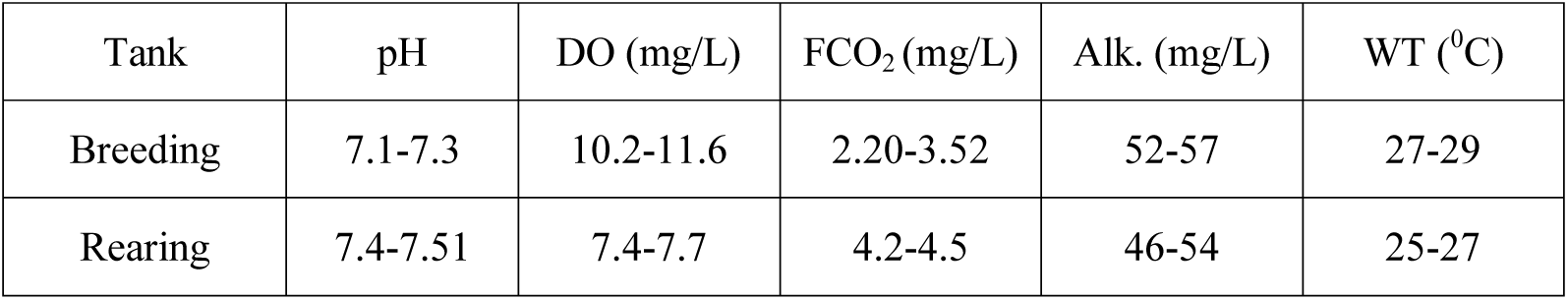
Physico-chemical parameter of water tank

| Tank | pH | DO (mg/L) | FCO<sub>2</sub> (mg/L) | Alk. (mg/L) | WT (°C) |
| --- | --- | --- | --- | --- | --- |
| Breeding | 7.1-7.3 | 10.2-11.6 | 2.20-3.52 | 52-57 | 27-29 |
| Rearing | 7.4-7.51 | 7.4-7.7 | 4.2-4.5 | 46-54 | 25-27 |

In the present study, *M. dibrugarensis* also showed more or less similar pattern of survival rate like as other *Mystus* species. Pond plankton and boiled chicken egg yolk mixed with water were provided to the hatchlings from 2^nd^ week onwards. After 15 days they were fed with finely macerated dried *Tubifex* when they were about three weeks. After 30 day, they were provided with chopped dried *Tubifex*, prawn, earthworm, insect larvae etc. The average rate of survival at different period was presented in Table 3.

**Table 3:** Survival and growth rate of hatchlings

| Age of fish | 5<sup>th</sup> day | 10<sup>th</sup> day | 15<sup>th</sup> day | 20<sup>th</sup> day | 30<sup>th</sup> day | Fingerlings |
| --- | --- | --- | --- | --- | --- | --- |
| Stocking density (no.) | 100 | 55 | 23 | 16 | 14 | 14 |
| Survivability (%) | 55 | 41.8 | 69 | 87 | 100 | 86 |
| Length (mm) | 6.2±0.81 | 11±3.31 | 12.1±1.05 | 15.3±3.07 | 21.1±2.04 | 49±3.03 |

## CONCLUSION

Fish in captivity may not always reproduce at the most favorable time. In this situation, proper dose of hormones play a critical role in the reproductive processes (Singh and Gupta, 2011). In the present study, was also found to breed naturally during April-July. Identical breeding period was also reported for *M. vittatus* by Arockiaraj *et al.* (2004) and Basu *et al.* (2015) and also in *M. tengara* (Rastogi and Saxsena, 1968). Mukherjee *et al.* (2002) able to breed *M. gulio* with ovaprim at dose of 2.5 ml/kg and at 0.4 ml/kg dose of ‘ovatide’ in *M. armatus*. Sarkar *et al.* (2005) reported induced spawning of *O. pabda* with single dose of ovaprim at 0.5 ml/kg resulting in 84-91 *%* fertilization rate and 65-66% hatching rate. Alam *et al.* (2006) found fertilization rate of 81-85% and hatching of 71-73% suggested that *M. gulio* also can successfully be bred in captivity using a single dose of 1-2 ml/kg body weight of both male and female.

Therefore, it may be concluded that in proper hormone dose, sex-ratio, and environmental condition the selected species can be successfully breed in captive condition. Their hardy nature, body design, wide food spectrum, acceptance of commercial pellets in captive conditions, ornamental value and high consumer preference for its taste, this endemic rarity ichthyofauna *M. dibrugarensis* make it a high value potential candidate for freshwater commercial aquaculture systems which is also an approach to conservation of this species.

## Acknowledgement

The authors gratefully acknowledge the Department of Life Sciences, Dibrugarh University for the laboratory facilities used during the study. Financial assistance from NFDB, Hyderabad is highly acknowledged.

---

**Fig 1A:**
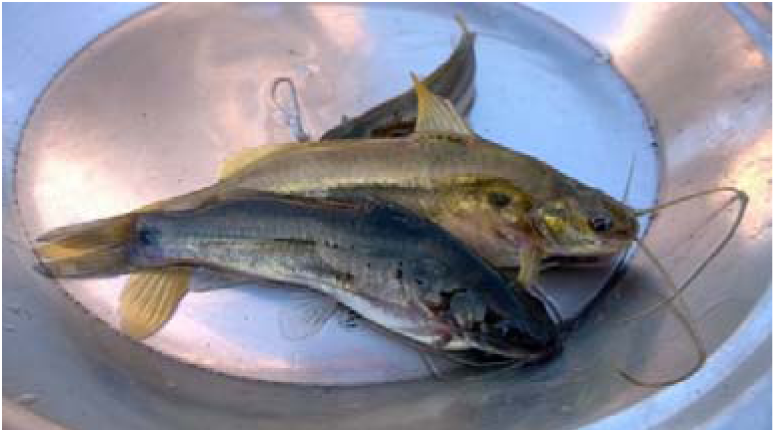
Female brooders of *M. dibrugarensis*.

**Fig 1B:**
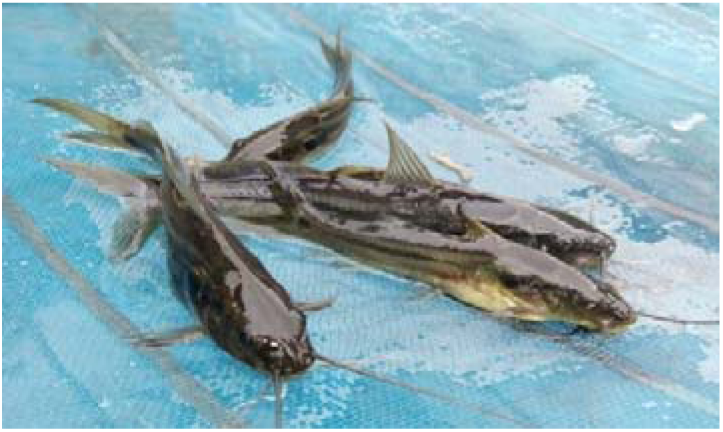
Male brooders.

**Fig 2:**
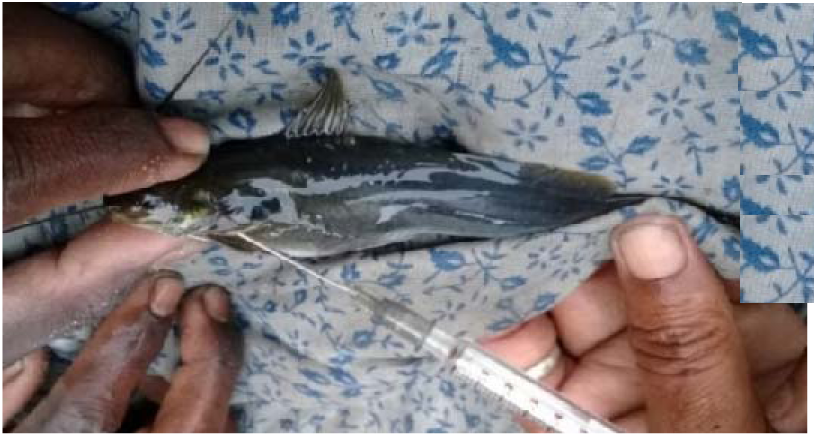
Hormone injection.

**Fig 3:**
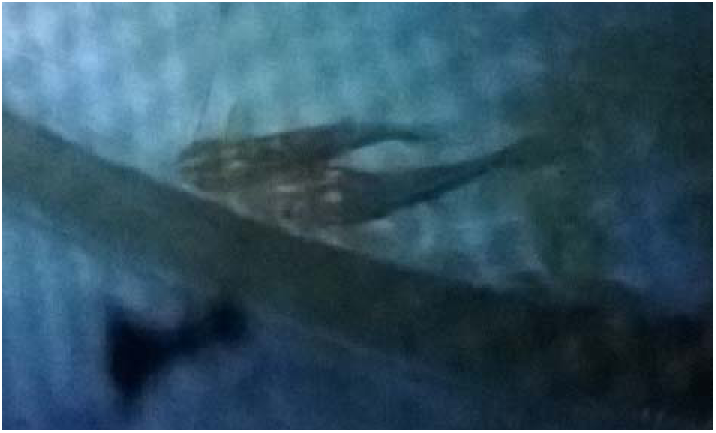
Pairing after injection.

**Fig 4:**
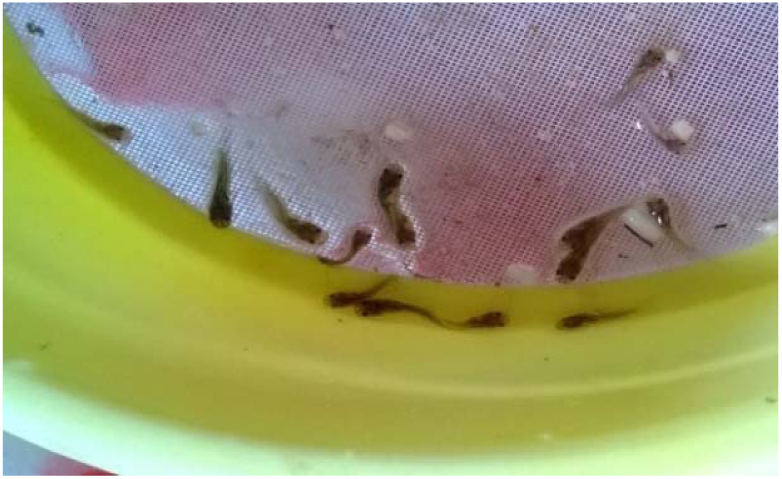
15^t^h day fry.

**Fig 5:**
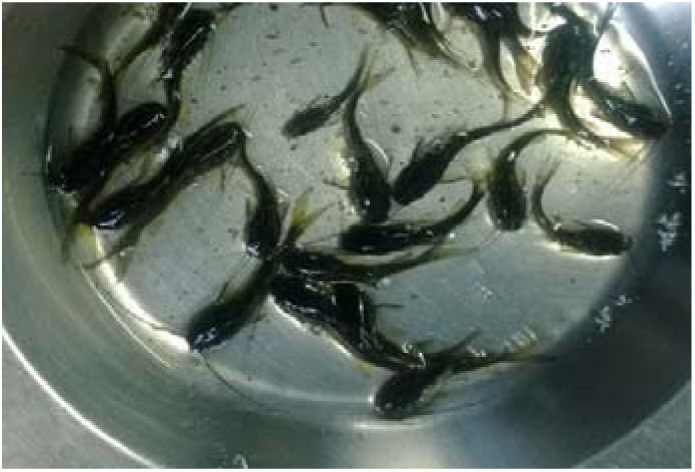
Fingerlings.

**Fig 6:**
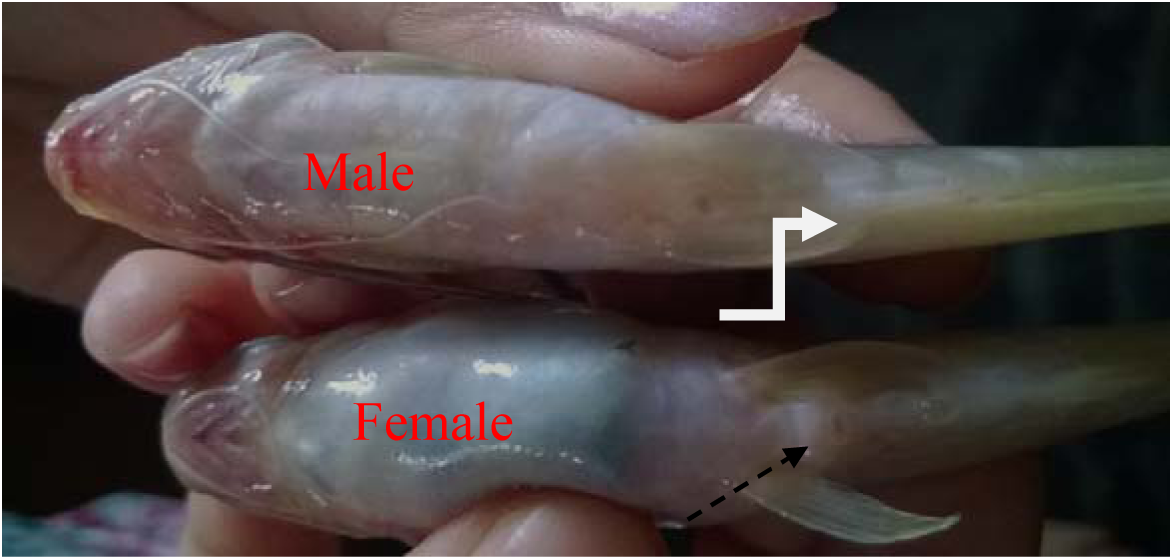
Sexual dimorphism in *M. dibrugarensis*.

